# A pooled-sample draft genome assembly provides insights into host plant-specific transcriptional responses of a Solanaceae-specializing pest, *Tupiocoris notatus* (Hemiptera: Miridae)

**DOI:** 10.1101/2023.10.27.564380

**Authors:** Jay K Goldberg, Carson W. Allan, Dario Copetti, Luciano M. Matzkin, Judith Bronstein

## Abstract

The assembly of genomes from pooled samples of genetically heterogenous samples of conspecifics remains challenging. In this study we show that high-quality genome assemblies can be produced from samples of multiple wild-caught individuals. We sequenced DNA extracted from a pooled sample of conspecific herbivorous insects (Hemiptera: Miridae: *Tupiocoris notatus*) acquired from a greenhouse infestation in Tucson, Arizona (in the range of 30-100 individuals; 0.5 mL tissue by volume) using PacBio highly accurate long reads (HiFi). The initial assembly contained multiple haplotigs (>85% BUSCOs duplicated), but duplicate contigs could be easily purged to reveal a highly complete assembly (95.6% BUSCO, 4.4% duplicated) that is highly contiguous by short-read assembly standards (N50 = 675 kb; Largest contig = 4.3 Mb). We then used our assembly as the basis for a genome-guided differential expression study of host-plant specific transcriptional responses. We found thousands of genes (N = 4982) to be differentially expressed between our new data from individuals feeding on *Datura wrightii* (Solanaceae) and existing RNA-seq data from *Nicotiana attenuata* (Solanaceae) fed individuals. We identified many of these genes as previously documented detoxification genes such as glutathione-S-transferases, cytochrome P450s, UDP-glucosyltransferases. Together our results show that long-read sequencing of pooled samples can provide a cost-effective genome assembly option for small insects and provide insights into the genetic mechanisms underlying interactions between plants and herbivorous pests.

## Introduction

Despite being highly toxic due to numerous noxious defensive chemicals, plants in the nightshade family (Solanaceae) have a handful of specialized herbivores. One of these insects is the tobacco suckfly (Hemiptera: Miridae: *Tupiocoris notatus*), which is known to feed on tobaccos (Nicotiana sp.; Halitschke et al. 2011) and sacred Datura (*Datura wrightii;* van Dam and Hare 1998), two genera known for their alkaloid defenses. *Tupiocoris notatus* has been an important part of research on the ecological roles of plant defense induction in wild tobacco (*Nicotiana attenuata*; Heidel and Baldwin 2004). Of note is *T. notatus*’ ability to ‘vaccinate’ plants against a more dangerous herbivore, the tobacco hornworm (*Manduca sexta* [Lepidoptera: Sphingidae]), a single individual of which can completely defoliate plants (Kessler and Baldwin 2004). It has also been implicated in the maintenance of a natural trichome dimorphism, by selecting against a glandular trichome producing morph when it becomes locally common (Goldberg et al. 2020). Furthermore, its transcriptome has previously been studied and putative plant-defense response genes have been identified (Crava et al. 2016). More recently, this species has been shown to manipulate plant defense and metabolism through cytokinins contained in its saliva (Brütting et al. 2018). Further insights into this insect and its ability to manipulate plant physiology for its own benefit would be vastly enabled with genomic resources that allow for deeper insights into the mechanistic underpinnings of its interactions with host plants. However, its small size presents difficulties for the generation of sequencing data with which to assemble a reference genome as it is not possible to extract enough DNA from a single individual for long-read sequencing platforms.

More generally, the assembly of genomes for small species of insect remains challenging due to problems associated with using pooled samples of individuals to generate sequencing reads. Past generations of DNA sequencing data (i.e. short reads and low-accuracy long reads) and assembly algorithms produce a single consensus sequence rather than phased haplotypes, and high heterozygosity can introduce errors to this process (Li et al. 2012). Newer sequencing data (i.e. PacBio HiFi reads and Oxford Nanopore Q20+ chemistry) and assembly algorithms are haplotype aware and designed to produce assemblies of separate haplotypes for single diploid organisms (Cheng et al. 2021). The presence of more than two haplotypes due to polyploidy or to the pooling of multiple individuals leads to the parallel assembly of multiple contigs (haplotigs) representing the same genomic region. Some species, such as many aphids, do not exhibit this problem due to the presence of an asexual stage in the life cycle, which produces a generation of genetically homogenous colonies in which multiple individuals can be pooled together without the risk of excess variation causing assembly errors (Davis 2012). Recently, bioinformatic solutions have been developed that allow for the removal of duplicated haplotigs from heterozygous genomes (Guan et al. 2020). However, their efficacy for removing duplicated contigs from pooled-sample assemblies has not yet been assessed.

In this study, we set out to produce a *T. notatus* draft genome assembly using sequencing data from a pooled sample of individuals. Using the purge_dups algorithm allowed us to generate a highly complete haploid genome assembly with most of the gene content represented, but with few allelic duplication errors. We further use this assembly as the basis for differential expression analyses to investigate its host-plant specific transcriptional responses.

## Methods

### Sample origins

In the fall of 2021, while growing *D. wrightii* for other studies, our greenhouse in Tucson, Arizona became infested with thousands of *T. notatus*. This small hemipteran herbivore specializes on plants with glandular trichomes, especially those in the Solanaceae (van Dam and Hare 1998). An originally small population likely entered the greenhouse without our knowledge sometime during the summer rainy season when insects are active in Southern Arizona and reached sufficient density to be noticed in the fall. Samples were collected via aspirator directly from *D. wrightii* plants in the greenhouse in December 2021 and January 2022. *Tupiocoris notatus* often co-occurs with a similar-looking predatory stiltbug (Hemiptera: Beritydae; Jay Goldberg *personal observations*), which we were careful to exclude during collections. All collections were immediately flash frozen with liquid nitrogen and stored at -80C.

### Nucleic acid extraction and sequencing

High molecular weight DNA was extracted from a single pooled sample of insects (0.5 mL, ∼50 individuals) using a previously established chloroform:isoamyl phase separation protocol (Jaworski et al. 2020). HMW DNA was size checked on by Femto Pulse System (Agilent) and 10 µg of DNA were sheared to appropriate size range (15-20 kb) using Megaruptor 3 (Diagenode). The sheared DNA was concentrated by bead purification using PB Beads (PacBio). The sequencing library was constructed following manufacturers protocols using SMRTbell Express Template Prep kit 2.0. The final library was size selected on a Pippin HT (Sage Science) using S1 marker with a 10-25 kb size selection. The recovered final library was quantified with Qubit HS kit (Invitrogen) and sized on Femto Pulse System (Agilent). The sequencing library was sequenced with PacBio Sequel II Sequencing kit 2.0, loaded to one 8M SMRT cell, and sequenced in CCS mode for 30 hours. RNA was extracted from similar pooled-samples (N = 3) using the ZYMO (Irvine, CA, USA) direct-zol miniprep kit (Cat. # R2050) and sequenced using NovaSeq (Illumina, San Diego, CA, USA) paired-end (150 bp) sequencing performed by Novogene (Sacramento, CA, USA).

### Genome assembly and annotation

CCS output (i.e.: HiFi reads; 3,583,689 reads; 17.69 Gb at mean Q35 score; mean length = 13,847 bp) were assembled using hifiasm-0.16.0 (Cheng et al. 2021). The initial assembly had numerous duplicated allelic contigs (Table 1, Figure 1) and was subjected to two rounds of the standalone purge_dups algorithm (v1.2.6; Guan et al. 2020). Assemblies were visualized using Bandage v0.8.1 (Wick et al. 2015), which also provided contiguity statistics. Contigs assembled from contaminant reads were identified and filtered from our assembly using the blobtools v1.1 pipeline (Laetsch and Blaxter 2017). Jellyfish v2.2.10 (Marcais and Kingford 2012) was used for k-mer counting before using the GenomeScope2.0 web portal (Ranallo-Benavidez et al. 2020) to estimate genome size (Figure S1). Filtered assembly quality and polishing was carried out using Inspector v1.0.2 (Chen et al. 2021). Gene content completeness was assessed via BUSCO v5.4.7 (Seppey et al. 2019) using the hemipteran odb10 dataset (Figure 1, Table 2). Repeat content of the final (twice purged, contaminant filtered, and polished) assembly was assessed using RepeatMasker v4.1.3 (Tarailo-Graovac and Chen 2009; Table S2) before structurally annotating gene content with the Helixer v0.3.1 algorithm pipeline (Stiehler et al. 2020; Holst et al. 2023) using the pre-made invertebrate training dataset. Functional annotation was done using InterProScan v5.45-80.0 (Jones et al. 2014) and blastp (using blast v2.13.0; Camacho et al. 2009) comparisons to the UniProt-Swissprot database (Boutet et al. 2007). Functional annotation outputs were combined into a single gff using the manage_functional_annotation.pl script in the AGAT v 1.2.0 toolkit (Dainat 2023).

**Table 1.** Summary statistics of assemblies before and after haplotig purging.

| **Hifiasm Assembly Statistics** |  |  |  |  |
| --- | --- | --- | --- | --- |
|  | Total length<br>(Mb) | Largest contig<br>(Mb) | N50 (kb) | # of<br>contigs |
| *Raw Assembly* | 1067.5 | 4.308 | 179 | 10061 |
| *Purged Once* | 405.7 | 4.308 | 542 | 1310 |
| *Purged Twice* | 296.1 | 4.308 | 665 | 908 |
| *Final Assembly* | 291.8 | 4.308 | 675 | 886 |

**Table 2.** BUSCO scoring results of assemblies and final annotation produced by Helixer.

| **BUSCO Scores (odb10 hemiptera; 2510 total genes)** |  |  |  |  |  |  |
| --- | --- | --- | --- | --- | --- | --- |
|  | Complete -<br>total | Single<br>Copy | Duplicated<br>(N ≥ 2) | Multi-<br>duplicated<br>(N ≥ 3) | Fragmented | Missing |
| *Raw Assembly* | 2440<br>(97.2%) | 254<br>(10.1%) | 2186<br>(87.1%) | 1658<br>(66.1%) | 22 (0.9%) | 48 (1.9%) |
| *Purged Once* | 2424<br>(96.6%) | 1569<br>(62.5%) | 855<br>(34.1%) | 112 (4.5%) | 29 (1.2%) | 57 (2.2%) |
| *Purged Twice* | 2416<br>(96.3%) | 2308<br>(92%) | 108 (4.3%) | 10 (0.44%) | 32 (1.3%) | 62 (2.4%) |
| *Final Assembly* | 2398<br>(95.6%) | 2288<br>(91.2%) | 110 (4.4%) | 11 (0.40%) | 30 (1.2%) | 82 (3.2%) |
| *Helixer*<br>*Annotation* | 2365<br>(94.2%) | 2244<br>(89.4%) | 121 (4.8%) | 9 (0.36%) | 47 (1.9%) | 98 (3.9%) |

**Figure 1.**
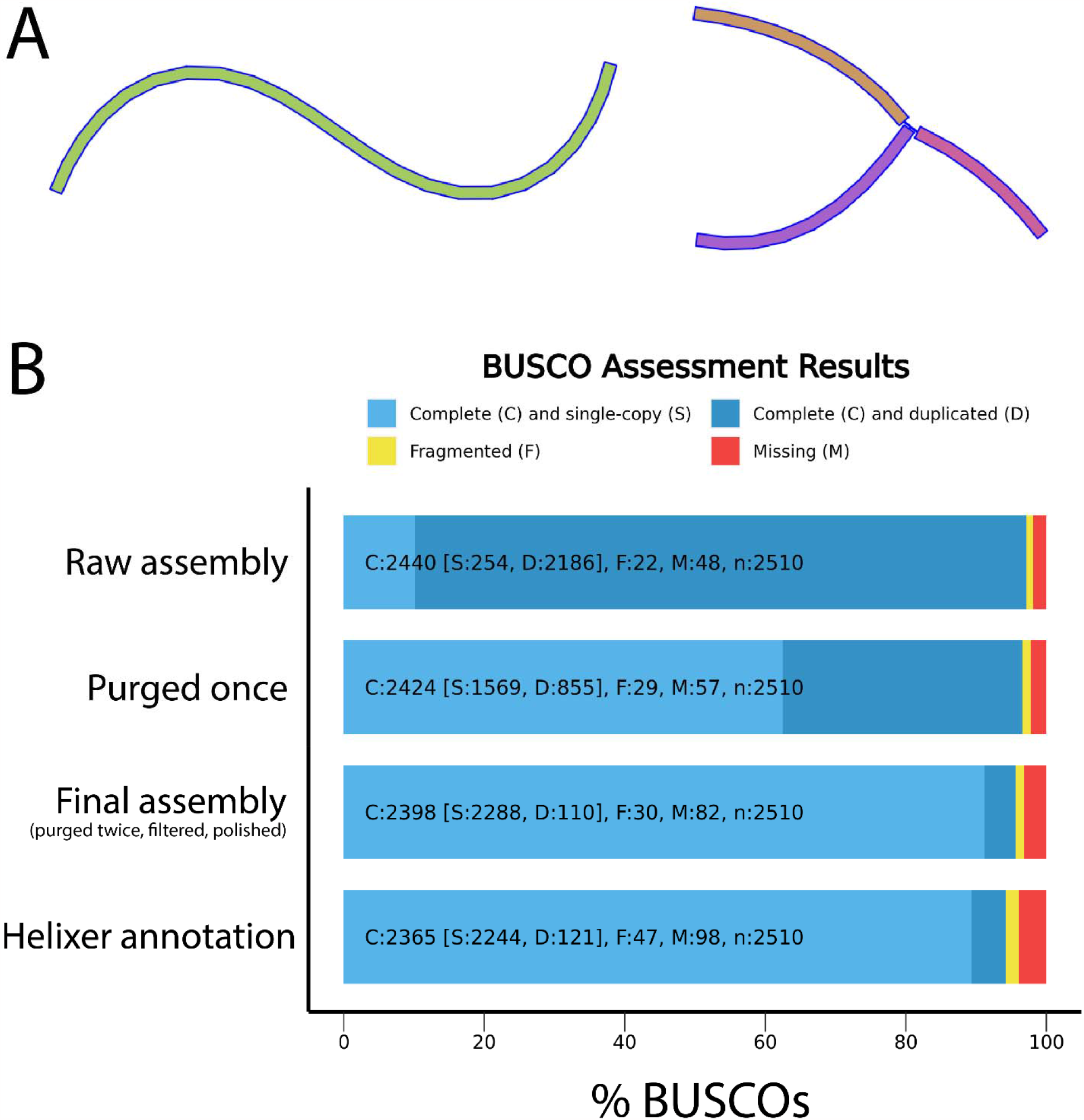
(A) Example of a properly assembled contig (left) and a y-shaped contig – indicative of unresolved repetitive elements – as found in the ‘raw’ assembly graphs (right). No y-shaped contigs were present in the final assembly. (B) Bar graphs showing the results of odb10 Hemiptera BUSCO analysis for the three of the genome assemblies we produced and our structural annotation of the final assembly (bottom bar). Annotation assessment was conducted in proteome mode.

### Differential expression analysis

RNA-seq reads were aligned to our genome assembly and counted using STAR (Dobin et al. 2013) using default settings after being trimmed with trimmomatic v0.39 (Bolger et al. 2014). Read counts were then analyzed using the DESeq2 package in R (Love et al. 2014; R Core Team 2021). We re-analyzed the existing dataset of tobacco-fed *T. notatus* transcriptomes (Crava et al. 2016; BioProject: PRJNA343704) and used all samples from their study as the baseline/control group (N = 6) for comparison to our *Datura wrightii*-fed dataset (N = 3). The ClusterProfiler v4.0 package was used for gene set enrichment analysis (GSEA; Wu et al. 2021) of differentially expressed genes. GSEA was performed on the total set of genes for which InterProScan obtained GO-terms. Each GO ontology (biological processes, molecular functions, and cellular component) was analyzed separately. We further separated each ontology into separate up- and down-regulated gene lists as prior studies have found this approach to be more robust than grouping all differentially expressed genes (DEGs) together (Hong et al. 2014). We used a significance cutoff of Padj=0.05 for all gene-wise analyses without any fold-change cutoff for differential expression.

## Results

### Genome assembly and annotation

The initial assembly was over 1 Gb in length (Table 1), making it highly duplicated (Fig 1) and far greater than the predicted size of 247 Mb. The first round of supplemental purging reduced this to 405 Mb (Table 1, Fig 1B), but left over 30% duplicated single-copy orthologues (Fig 1B). A second round of purge_dups reduced this to a reasonable level (Table 1, 2). The size of the twice purged assembly (296 Mb, Table 1) was closer to the predicted size (247 Mb, Fig S1). Structural annotation identified 16,067 genes and the protein dataset had 95.1% of BUSCO genes complete when compared to the hemiptera_odb10 reference dataset (Fig 1B; Table 2). Taxonomic identification via comparison to the nt database (Camacho et al. 2009) in blobtools found 22 low-coverage contigs likely to originate from contaminant reads (Figure S2; Table 1). These were filtered out of our assembly before beginning downstream analyses. Quality assessment with Inspector yielded an initial QV-score of 19.7, roughly 1500 structural errors, and a small-scale error rate of 4132.5 errors per Mb. After polishing the QV-score increased to 21.7 and the small-scale error-rate was reduced to 116 per Mb. Structural errors were largely unchanged by the polishing process and slightly increased from 1513 to 1542. Nearly all raw reads (99%) of raw reads mapped back to the polished assembly for an average read depth of 60.1. Detailed output of Inspector analyses pre- and post-polishing are in Table S1. RepeatMasker found 34.98% of the final assembly to be composed of repetitive elements (6.35% retroelements, 1.09% DNA transposons, 2.09% rolling-circle transposons, 24.27% unclassified; detailed output in Table S2). Helixer annotated 16062 genes in our final assembly, ranging in size from 108 bp to 379 kb (mean = 11.9 kb). 13875 of these genes were functionally annotated, including 8824 for which GO-terms could be assigned. Detailed annotation statistics are found in Table S3. Overall, these results show that the quality of our assembly lags behind that of recent chromosome-scale assemblies produced using combined long-read sequencing and chromatin conformation capture technologies (e.g. Hi-C; Wang et al. 2023) in terms of accuracy and contiguity.

### Differential expression analysis

Alignment of RNA-seq reads was consistent across samples. An average of 94.8% of raw reads were mapped to our assembly (min = 92.1%, max = 96.1%). Most reads (mean = 77.7%) mapped uniquely, but many (mean = 17.1%) mapped to multiple loci. Few reads (mean = 0.6%) were thrown out due to excessive multi-mapping. Most reads that were unused were too short for mapping (mean = 4.53%). No reads were found to have too many mismatches to be used. Full read-mapping statistics can be found in Table S4. Only a small proportion of our annotated genes (N = 509) did not have any mapped reads.

The dataset used by Crava et al. (2016) used a de novo transcriptome approach to look at differentially expressed genes in *T. notatus* feeding on a transgenic line of wild tobacco (*Nicotiana attenuata*). The ability to produce jasmonic acid and induce defense production is compromised via RNAi silencing of the biosynthetic gene allene oxide cyclase (irAOC, N = 3; vs empty vector controls, EV; N = 3). We reanalyzed their dataset using a genome-guided approach enabled by our draft assembly (Figure 2, Table S5). Our principal component analysis (PCA) found that iAOC and EV fed insects formed separate clusters (Figure 2A) indicating different patterns of gene expression in each sample set; however, when we examined these patterns gene-by-gene, we only found 11 significantly differentially expressed loci (8 down-regulated, 3 up-regulated; Figure 2B). None of these genes showed substantial levels of expression changes (Log_2_fold-change < 1). Six of these genes were functionally annotated (Table S5), but none belonged to the detoxification gene families (cytochrome P450s, glutathione-S-transferases, UDP-glucuronosyltransferases) identified in their study (Crava et al. 2016). One down-regulated gene was associated with digestion of plant-compounds and annotated as polygalacturonase (PGN1). The difference between our results could be due to the presence of split or non-coding genes due to Trinity assembly errors (Grabherr et al. 2011; Freedman et al. 2020) in their de novo transcriptome dataset, which contained 42610 putative genes; a far larger number that was annotated within our assembly. It is also possible that the difference is an artifact of reference-derived biases, as our genome was assembled from individuals originating in the Tucson (Arizona) area and their RNA-seq data was collected from a population in Utah. The genetic variation of this species is not known and it is possible that presence-absence variation in gene content exists – should there be substantial population structuring – as it does in other herbivorous insects (Mongue and Kawahara 2022). This is likely to only be the case if there is geographic variation in expression levels, as we did not observe substantial differences in mapping rates between the two datasets (*N. attenuata*, Mean = 95.15%; *D. wrightii*, Mean = 94.24%).

**Figure 2.**
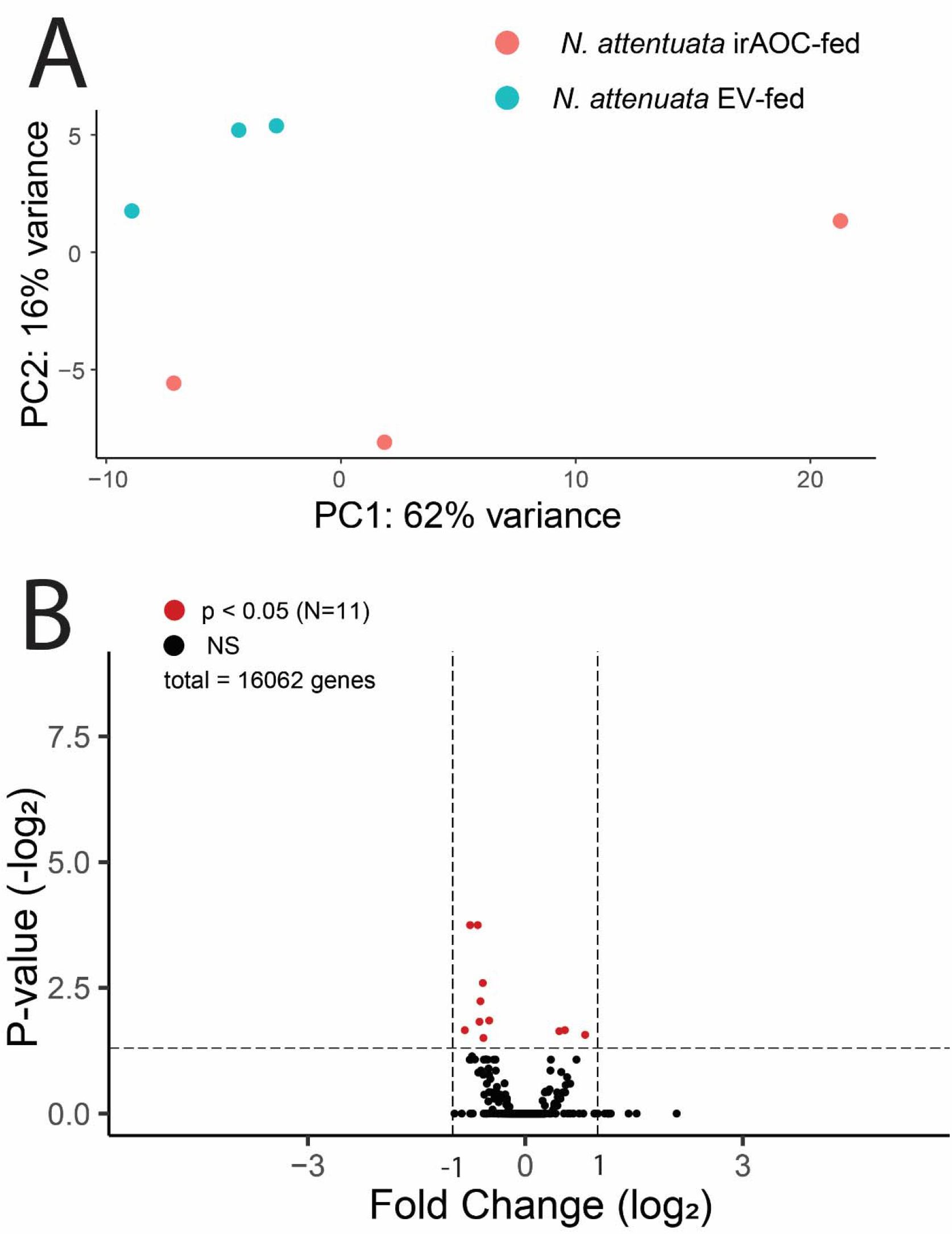
(A) Results of PCA analysis from differential expression study that included only Nicotiana-fed insects from a previous study (Crava et al 2016). (A) PCA analysis shows that the two treatments do not form distinct clusters and are not substantially different when gene expression is viewed globally. (B) Volcano plot showing the p-value associated with the ‘line’ variable in differential expression analysis and the fold change of that gene. Two-fold changes (|log_2_| = 1) in either direction are marked for reference, but this was not used as a testing cutoff. Genes that satisfied our cutoff for significance (P_adj_ < 0.05) are shown in red (N = 50; Table S4), whereas NS genes are in black.

We found that the expression profiles of *Datura*- and *Nicotiana*-fed insects were distinct (Fig 3A). This difference was driven by many significantly up-or down-regulated genes (N_total_ = 4982; N_down_ = 2121; N_up_ = 2861; Fig 3B, Table S2). The most drastic differences in expression had over 10-fold changes in either direction (272 genes with |Log_2_FC| > 3.01). Six differentially expressed genes (DEGs) were functionally annotated as glutathione-S-transferases, all of which were found to be up-regulated in *Datura wrightii*-fed samples. Another seven DEGs were annotated as UDP-glycosyltransferases, four of which were down-regulated and the other three up-regulated in *Datura*-fed insects. 45 cytochrome P450s were identified as DEG and most of them (N = 29) were up-regulated. Out of 22 differentially expressed serine proteases, 17 were found to be upregulated in our *Datura*-fed samples. The full list of significantly differentially expressed genes, including fold-change and adjusted p-values, is in Table S6. Our findings are consistent with previous studies of transcriptomic responses of herbivores to host-plant chemistry (Castañeda et al. 2009, Bock et al. 2016, Lin et al. 2021) and confirm the role of the aforementioned gene families in digestion/detoxification of plant-derived compounds by *T. notatus*.

**Figure 3.**
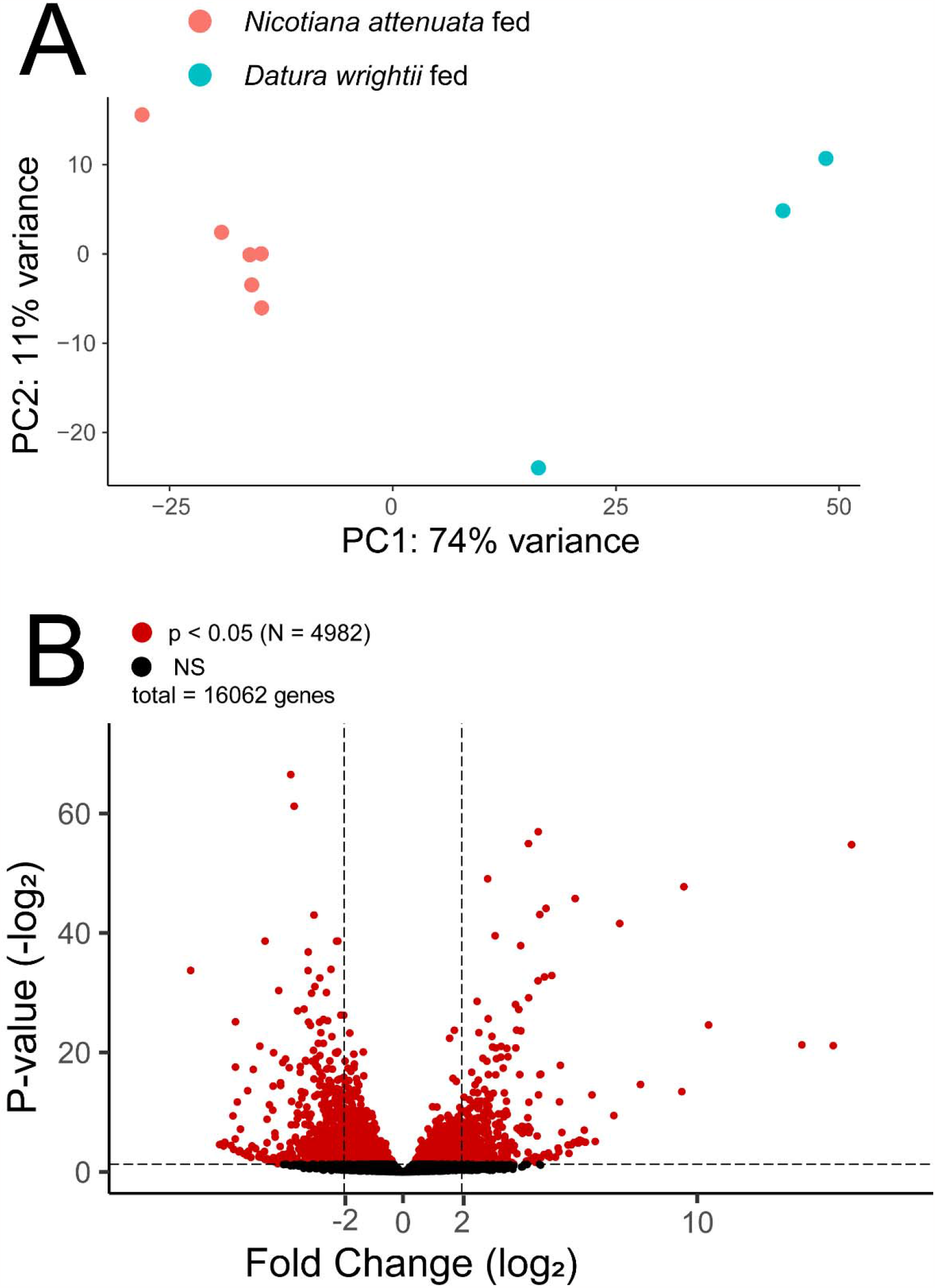
(A) Results of PCA analysis from differential expression study. *Nicotiana* and *Datura*-fed insects form distinct clusters. (B) Volcano plot showing the p-value associated with the ‘plant’ variable in differential expression analysis and the fold change of that gene. Four-fold changes in either direction (|log_2_| = 2) are marked for reference, but this was not used as a testing cutoff. Genes that satisfied our cutoff for significance (P_adj_ < 0.05) are shown in red (N = 4982; Table S3), whereas NS genes are in black.

To explore the transcriptional changes associated with host-plant species beyond our handful of target genes, we used a gene set enrichment analysis to identify common themes in our set of DEGs. We found a total of 24 GO terms to be significantly enriched within our results (Figure 4, Table S7) and that these are associated with gene/protein expression, nutrient catabolism, and chemosensory functions. Digestive functions were predominantly up-regulated, with the notable exception of aspartic-type endopeptidases. Another notable finding is that gustatory perception is enriched in both up- and down-regulated pathways, suggesting the presence of complex host-plant associated changes to chemosensory pathways. Olfaction was found to only be enriched within up-regulated pathways.

**Figure 4.**
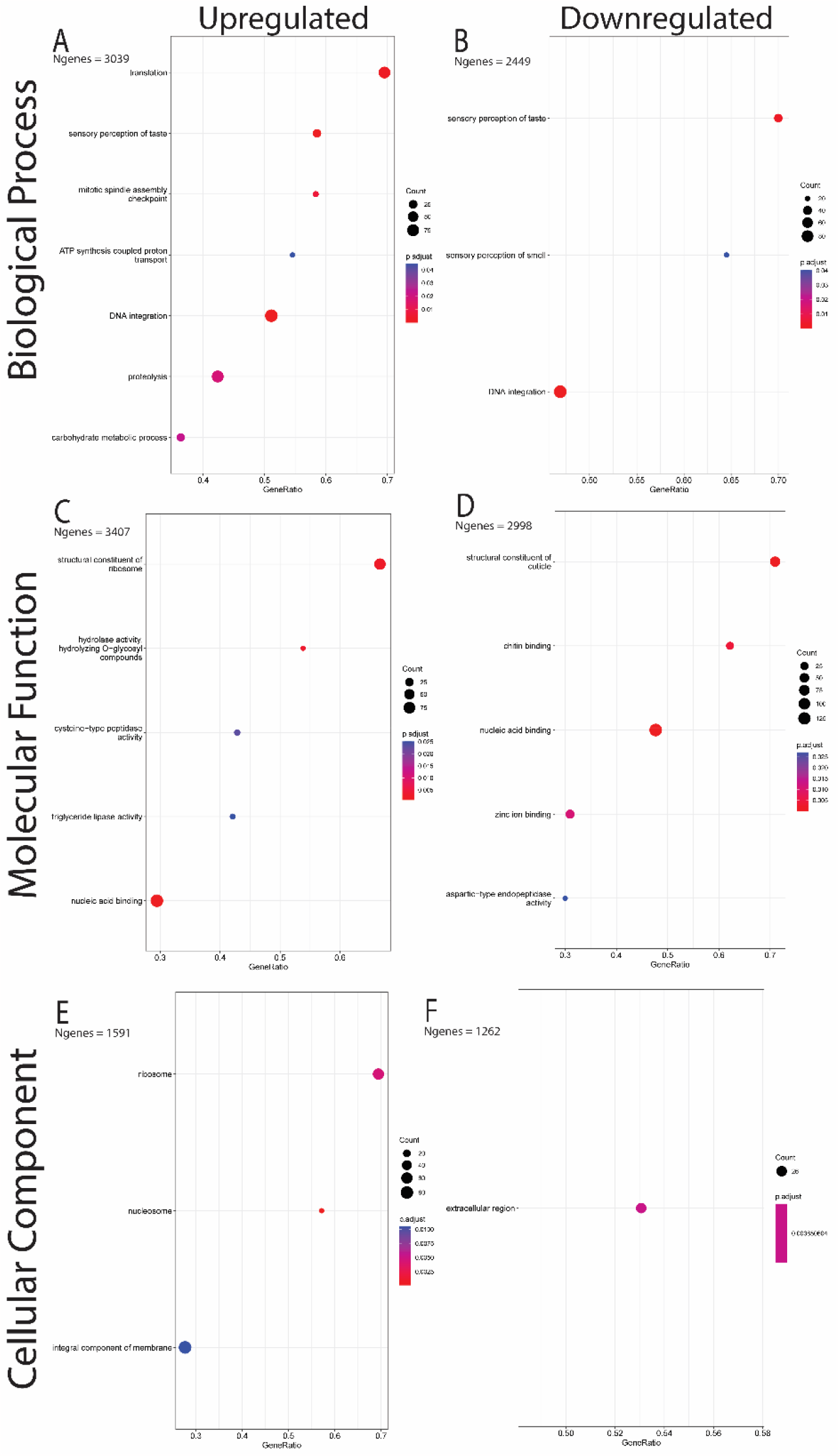
Results of gene set enrichment analyses. 24 Go terms were found to be enriched in either up-or down-regulated gene lists. X-axes show the ratio of enriched genes vs the total count of genes sharing that GO term. Dot sizes represent the total number of genes sharing each GO term whereas dot colors represent the p-value (adjusted for multiple tests) of each term.

## Discussion

Producing high-quality reference genome assemblies for small insects remains challenging. In this study we were able to produce a high-quality draft assembly using PacBio HiFi reads from a pooled-sample of genetically variable individuals. It is important to note that our assembly is not chromosome-scale or error-free, and should be not used as a reference in analyses such as studies of chromosomal rearrangements or other structural variants. However, the unique protein-coding regions are well-represented, indicating that our assembly is of sufficient quality for studies focused specifically on low-copy gene content and function. Moreover, our assembly was far more contiguous and complete than genome assemblies produced using short-read technologies. Repetitive element and gene content of our assembly is comparable to that of another mirid (*Cyrtorhinus lividipennis*) which was previously found to have a 345.75 Mb genome containing 14,644 protein-coding genes and 31.1% repeat content (Bai et al. 2021). In its current state, it is a suitable reference for the gene content of this species. We were able to use this genome as the basis for multiple differential expression analyses and preliminary assessments of functional gene content. It will likely also serve as a suitable reference assembly for population genomic analyses and other read-mapping based pipelines. This demonstrates the utility of pooled-sample genome assemblies when working with small insects that present difficulties for extracting sufficient quantities of DNA from a single individual.

The first of our differential expression studies was a re-analysis of a previously published dataset comparing wild tobacco plants (*N. attenuata*) with functional (empty vector control; EV) and compromised (via RNAi silencing of allene oxide synthase expression; irAOC) defense induction pathways. We found that few insect genes were differentially expressed between colonies fed on these two lines, and that none of the significant genes were strongly up-or down-regulated. This indicates that jasmonate-induced plant defenses may not play a role in interactions between plants and *T. notatus*; a stark contrast to interactions between *N. attenuata* and other herbivores, such as *M. sexta*, against which induced defenses have been shown to play a critical role (van Dam et al. 2000). This finding is also different from the study that generated these RNA-seq data, which found dozens of significantly differentially expressed genes (Crava et al. 2016) and is likely due to the more conservative nature of our genome-guided approach compared to the de novo transcriptome assembly used as a mapping reference in their analysis.

In addition to our reanalysis of Crava et al (2016)’s dataset, we also compared newly generated RNA-Seq data from our *D. wrightii*-fed greenhouse population to their *N. attenuata*-fed data. Our pooled samples were collected and prepared in a similar fashion to theirs, although differences may be present due to the geography of the sampled populations as little is known about population structure in this species. Their data originated from a captive colony (maintained in Jena, Germany) started from individuals collected from a field site in Southwest Utah. Our samples were collected from a greenhouse infestation in Tucson, Arizona roughly 615 km away from their field site. Nonetheless we identified many detoxification, digestion, and chemosensory genes in our list of differentially expressed genes. This finding suggests that a substantial amount of the gene expression differences between our samples and Crava et al. (2016)’s is due to host-plant specific responses. We consider our list of genes a suitable starting point for future studies into the genetic basis of interactions between this species and its toxic Solanaceous hosts. Future studies might examine expression differences in response to more controlled manipulation of specific plant defensive compounds, or tissue-specific gene expression by *T. notatus*, to differentiate between genes involved with plant-metabolism manipulation – which are likely to be expressed in salivary glands (Boulain et al. 2019) – from those involved with digestion/detoxification. Overall, our results suggest the presence of physiological differences between *T. notatus* feeding on *Datura* and *Nicotiana*. Many of these differences are related to perception, digestion, and detoxification of host-plant derived compounds, but others could also be derived from population (Arizona vs. Utah) differences or responses to other factors (e.g. greenhouse conditions). Our findings nonetheless provide a valuable starting point for future targeted studies of differentially expressed genes with specific roles mediating host-plant interactions.

In conclusion, we have demonstrated that by using a standard haplotig purging algorithm, high-quality pooled-sample genome assemblies of a single haplotype can be produced. We have demonstrated the utility of our assembly for RNA-Seq read-mapping based pipelines by conducting two genome-guided differential expression studies. We identified differentially expressed genes associated with specific host-plant interactions and provide an initial functional assessment of them, many of which belong to well-known families of detoxification and digestion genes. Together, these findings show that pooled samples may be a viable option for researchers unable to sequence single individuals of their species of interest due to small size or other factors and provide a valuable starting point for future research into the interactions between specialist herbivores (Hemiptera: Miridae) and their host plants.

## Supporting information

Table S1 - Inspector Results

Table S2 - RepeatMasker Results

Table S3 - Annotation Statistics from AGAT

Table S4 - RNAseq read mapping statistics

Table S5 - Nicotiana attenuata only resutls

Table S6 - Datura vs Nicotiana results

Table S7 - GSEA results

High resolution version of Figure 4 for enchanced readability

## Data Availability

Sequencing data, including raw reads and the final genome assembly, are deposited on NCBI under BioProject PRJNA971612 (reviewer link: https://dataview.ncbi.nlm.nih.gov/object/PRJNA971612?reviewer=vb457840npkmq59gac1dm9k4b6). No new code or analyses were generated for this project, but shell and R scripts used to run existing programs/packages are located at https://github.com/caterpillar-coevolution/Tupiocoris-notatus-genome-project. Additional datasets not included as supplemental materials, such as the genome annotation files, can be found in the GitHub repository as well. Published data from Crava et al. (2016) are available under BioProject: PRJNA343704.

## Acknowledgements

Computational analyses were conducted on the UArizona HPC and we would like to thank the staff for maintaining and providing access to this resource. We would like to thank Alex Karnish and Beth-Ann Hansen for taking care of greenhouse plants and allowing the collection of insects before eliminating the *T. notatus* infestation.

## Author Contributions

The project was conceived of, and analyses conducted by JKG with substantial input and assistance from CWA, DC, LMM, and JB. JKG drafted the manuscript and all authors reviewed and contributed to the final version of the manuscript.

