## Supplemental Figures & Table Legends for "A pooled-sample draft genome assembly provides insights into host plant-specific transcriptional responses of a Solanaceae-specializing pest, *Tupiocoris notatus* (Hemiptera: Miridae)"

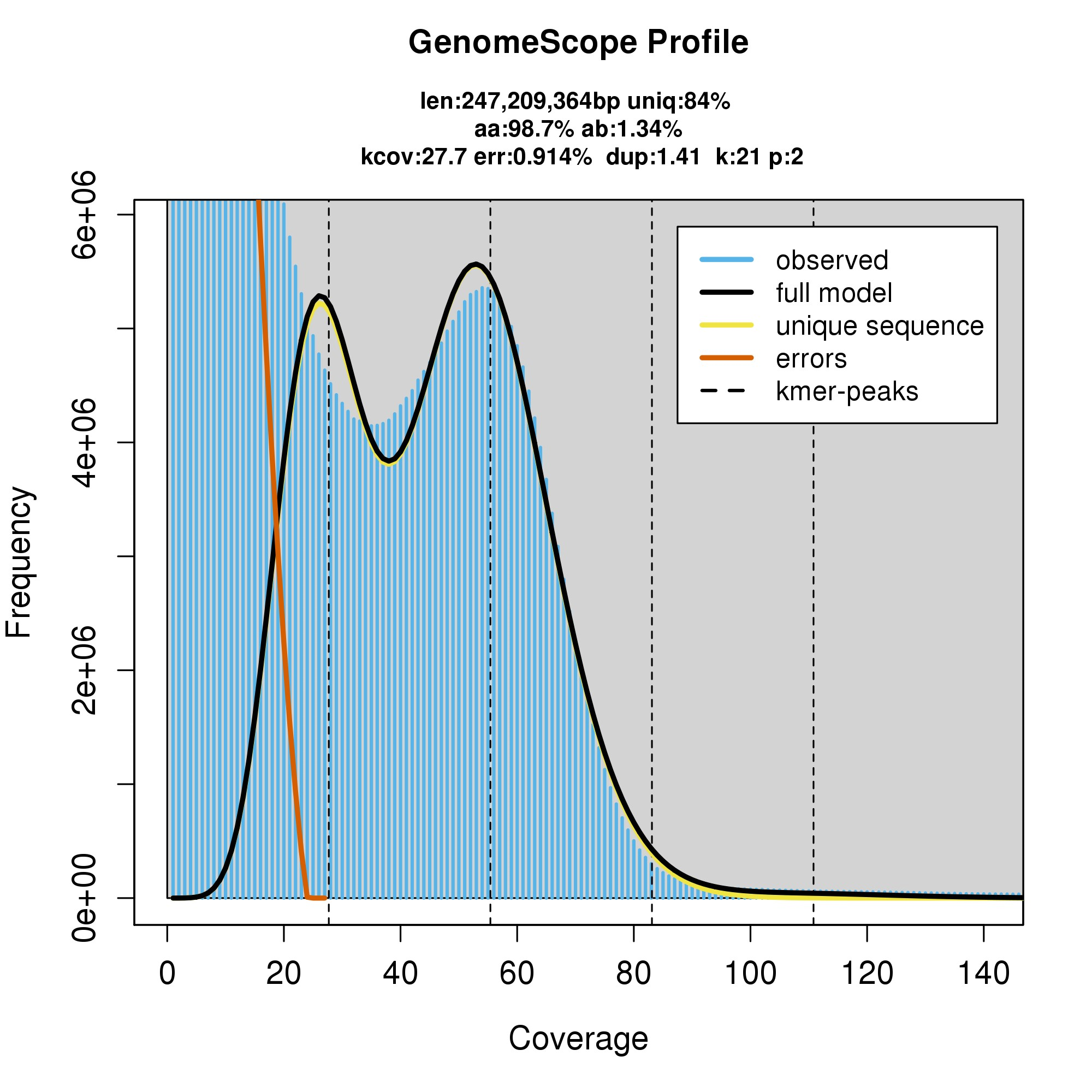

**Figure S1.** K-mer coverage plot generated by GenomeScope. Our dataset had a highly elevated frequency of low coverage k-mers due to the variation introduced by sequencing a pooled sample of individuals, however, the frequency of homozygous alleles was nearly that predicted by the model.

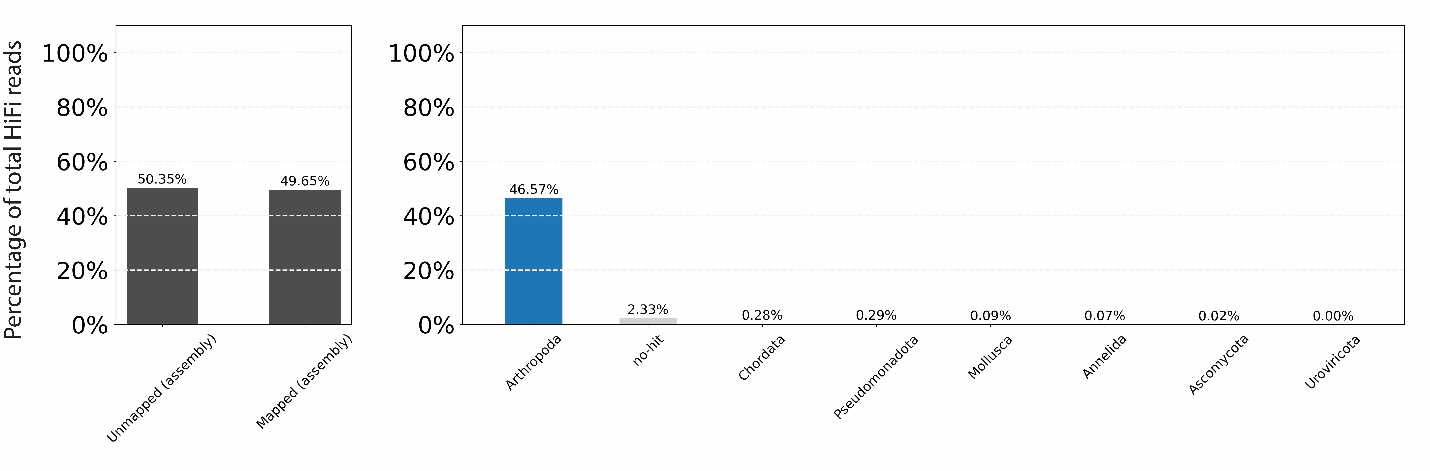

**Figure S2.** Coverage plots produced by the blobtools pipeline. **(left)** shows the percentage of raw reads that mapped back to the final (twice purged) assembly. **(right)** shows the percentage (of the total raw reads) that mapped to contigs with the associated taxaID. Most contigs were associated with phylum Arthropoda. Contigs that were identified as other phyla were filtered from the assembly. No-hit contigs were kept in our final assembly.

**Table S1.** Inspector output with assembly statistics before and after polishing contigs

**Table S2.** RepeatMasker output describing repetitive element content

**Table S3.** AGAT output with detailed structural annotation statistics

**Table S4.** STAR output regarding the alignment of RNA-seq reads

**Table S5.** Complete list of significant DEGs from reanalysis of NaEV vs irAOC-fed insects

**Table S6.** Complete list of significant DEGs in Datura- vs tobacco-fed insects

**Table S7.** Complete list of enrichment analysis and summary statistics
