## Supplementary figures and images for "A pooled-sample draft genome assembly provides insights into host plant-specific transcriptional responses of a Solanaceae-specializing pest, *Tupiocoris notatus* (Hemiptera: Miridae)"

### High resolution version of Figure 4 for enchanced readability

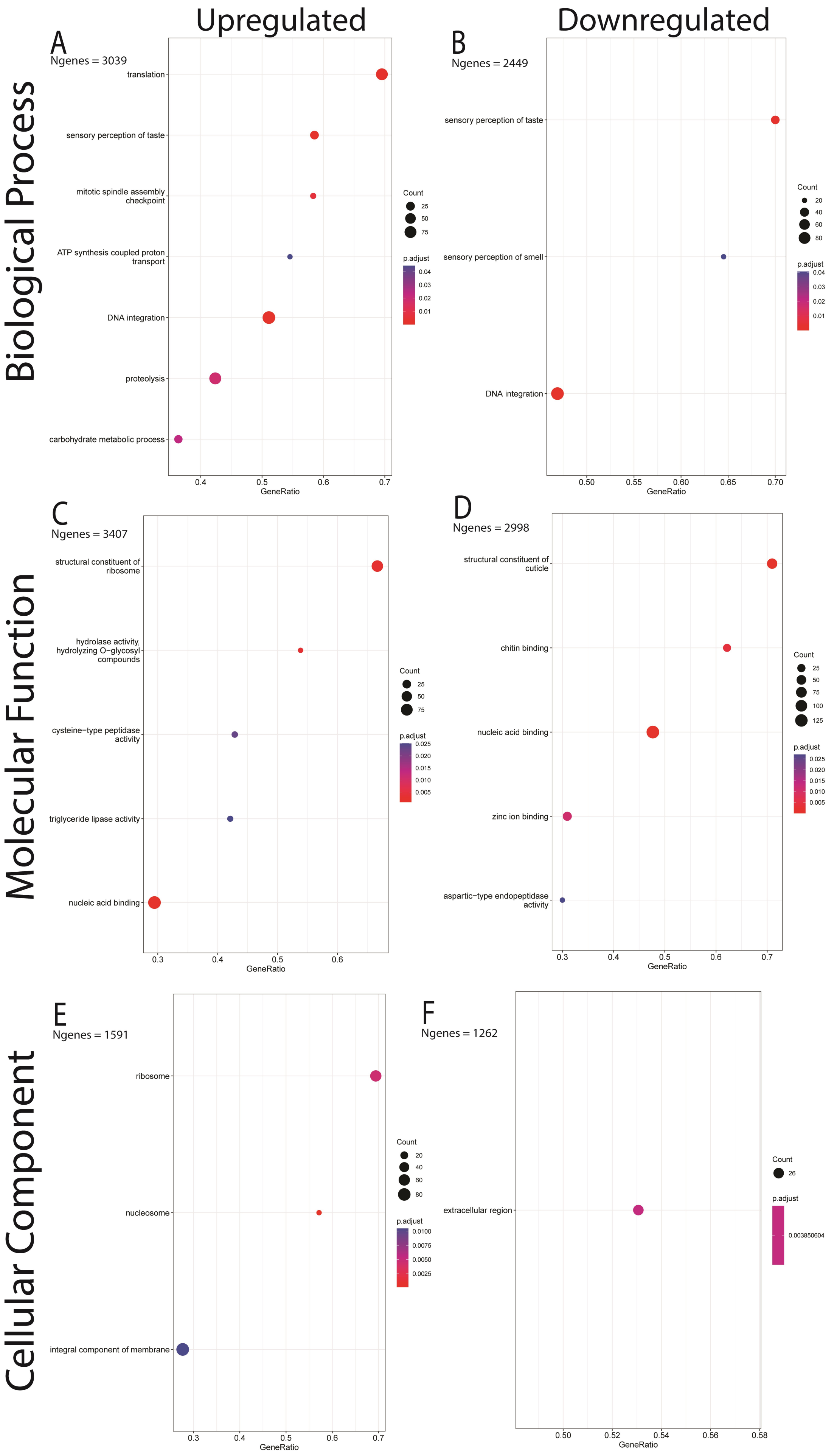
